# Cell Contractile Force-Mediated Morphodynamical Tissue Engineering via 4D Printed Degradable Hydrogel Scaffolds

**DOI:** 10.1101/2025.02.22.639682

**Authors:** Aixiang Ding, Kaelyn L. Gasvoda, David S. Cleveland, Eben Alsberg

## Abstract

Tissue morphogenesis is a critical aspect tissue development. Recent advances in four-dimensional (4D) cell scaffolds have shown promise for modeling morphogenic processes. While current 4D systems often rely on external stimuli, they tend to overlook the role of intrinsic cell-generated forces, such as cell contractile forces (CCFs), in driving tissue morphogenesis. The paradox between the inherent weakness of CCFs and the robustness of tissue scaffolds presents a significant challenge in achieving effective shape transformations. In this study, we introduce an easily printable, freestanding, cell-laden hydrogel platform designed to harness CCFs for 4D shape morphing. These hydrogels initially provide mechanical support to maintain structural integrity, followed by rapid degradation that amplifies CCFs through enhanced cell-cell interactions and increased local cell density, thereby inducing tissue morphogenesis. This platform enables the formation of scaffold-free constructs with programmed shape transformations. By modulating the initial printed geometries, complex and large tissue constructs can be generated via controlled global shape transformations. Furthermore, the platform supports 4D tissue engineering by facilitating tissue differentiation coupled with dynamic shape evolution. This CCF-4D system represents a significant advancement in biomimetic tissue engineering, offering new avenues for creating dynamic tissue models that closely replicate native morphogenesis.

## Introduction

Tissue morphogenesis, a critical aspect of biological development, has been widely studied due to its fundamental role in the formation of functional tissue structures^1-3^. This process involves the organization of multicellular tissues into specific morphological architectures, driven by complex cellular interactions, including cell-cell and cell-extracellular matrix (ECM) interactions^4^. Such interactions are essential for achieving and maintaining tissue function, making the study of tissue morphogenesis highly relevant within the field of tissue engineering. Tissue engineering platforms provide a means to mimic and study tissue development *in vitro*^5,6^, using models such as cell condensations^6^ and hydrogel matrices^7,8^ to replicate tissue morphogenesis.

Cell condensation models are typically characterized by cell aggregation with initially minimal ECM components, forming tightly packed structures known as spheroids^9^. Cell spheroids, characterized by strong cellular interactions^6^ and rapid formation^9,10^, are frequently utilized for morphogenic studies. However, these models have limited capacity to organize into more complex hierarchical structures, thereby restricting their ability to undergo dynamic morphogenic processes such as bending, curling, folding, and buckling, which are critical to tissue development *in vivo*^2-4^. Conversely, conventional cell-laden hydrogel models^11-13^, while providing a more defined structure, often result in rigid constructs that limit cell-cell interactions and cellular rearrangement. These limitations hinder the replication of the dynamic curvature patterns observed in native tissue morphogenesis. Given these constraints, there is a need for alternative modeling strategies that better replicate the dynamic processes of tissue morphogenesis observed in vivo, including the complex curvature patterns formed through cell-driven folding and bending.

Four-dimensional (4D) biofabrication, an emerging approach in tissue engineering, incorporates time as an additional dimension, allowing for dynamic changes in scaffold shape to better mimic biological processes^13-19^. These geometric transformations were first observed in biological development and healing^10^ and have since been applied in various studies^14-17,20,21^. Methods for inducing shape changes in 4D scaffolds typically involve the fabrication of multi-material systems with differential swelling^14,15,22,23^ and employs various stimuli, including infrared light^24,25^, temperature variations^26,27^, electrical^28^ or magnetic fields^29^, and chemical additives^30^. However, these methods often rely on external stimuli, neglecting the intrinsic cellular forces that can play a role in driving tissue morphogenesis^31,32^, which limits their ability to partially replicate this aspect of in vivo tissue formation processes ^31^. Consequently, it may be valuable to develop more sophisticated 4D scaffolds that leverage endogenous biological mechanisms to replicate the dynamic changes of tissue architectures seen in living organisms.

Recent research has highlighted the critical role of cell contractile force (CCF) in driving complex tissue architecture formation in vivo^32,33^. Leveraging CCF to induce scaffold deformation presents a promising strategy for controlling 4D shape changes with potential clinical applications. Previous studies have explored CCF-driven shape changes using engineered scaffolds, such as microplates^34^, DNA Velcro^35,36^, and photolithography-assisted extrusion bioprinting^37^. However, these methods face limitations in scalability, shape morphing resolution, and applicability in clinical settings due to challenges in achieving stable, large-scale constructs and replicating the dynamic morphological changes required for in vivo-like tissue formation. For example, 4D microplate structures that rely on cell seeding onto non-biodegradable substrates encounter challenges in replicating the native three-dimensional (3D) environments, where cells are embedded within the surrounding tissue matrices^34^. DNA Velcro, another approach for achieving CCF-mediated 4D shape changes, involves the deposition of loose cell aggregates onto superficial regions of a substrate^35,36^. Although this method can successfully produce specific geometric changes, its reliance on complex procedures for precisely controlling the spatial positioning of cell aggregates presents challenge for high throughput implementation. Extrusion bioprinting has been explored as a potential strategy for engineering large-scale CCF-based 4D tissue constructs with patterned cell distributions^37^. However, the reported approach faces challenges related to limited shape-morphing resolution over short timeframes, likely due to the softness of the materials, which makes it difficult to print stable, freestanding constructs. Consequently, current CCF-driven systems are inadequate for achieving complex shape morphing on a large scale or replicating the dynamic 4D tissue formation observed in vivo. Additionally, the lack of gradual or controlled degradation of the scaffolds used further restricts their use in clinical applications and in vivo studies, as it impedes proper cell organization, rearrangement, and remodeling within the system. To address these challenges, there is a need to develop CCF-driven actuation systems capable of generating stable, large tissue constructs without residual biomaterials that undergo 4D geometric shape changes to replicate the complex curvature patterns seen in native tissue morphogenesis.

In this study, we present a transformable, single-material scaffold composed of a rapidly degradable hydrogel support structure fabricated via 4D bioprinting, which harnesses the inherent CCF exerted by encapsulated cells to achieve dynamic geometric transformations that partially replicate aspects of *in vivo* tissue morphogenesis (**Scheme 1a**). This scaffold consists of oxidized methacrylated alginate (OMA) microgels and cell-adhesive gelatin methacrylate (GelMA), homogeneously embedded with live cells. Within this composite system, CCF drives scaffold deformation, leading to the formation of tissue-only (scaffold material-free) tissue constructs with predefined geometries following scaffold degradation and tissue maturation (**Scheme 1b**). This approach offers significant advantages over existing methods by: (1) enabling the formation of more complex shapes and morphing patterns due to the high printability of the microgel bioink, which allows for the creation of freestanding constructs with high resolution and fidelity, and (2) utilizing CCF as a universal morphogenic driver that supports long-term 4D tissue formation. Our system thus provides a more biomimetic platform for replicating the specific geometries and morphologies of tissue development in vivo, with potential applications in regenerative medicine and tissue engineering.

**Scheme 1.**
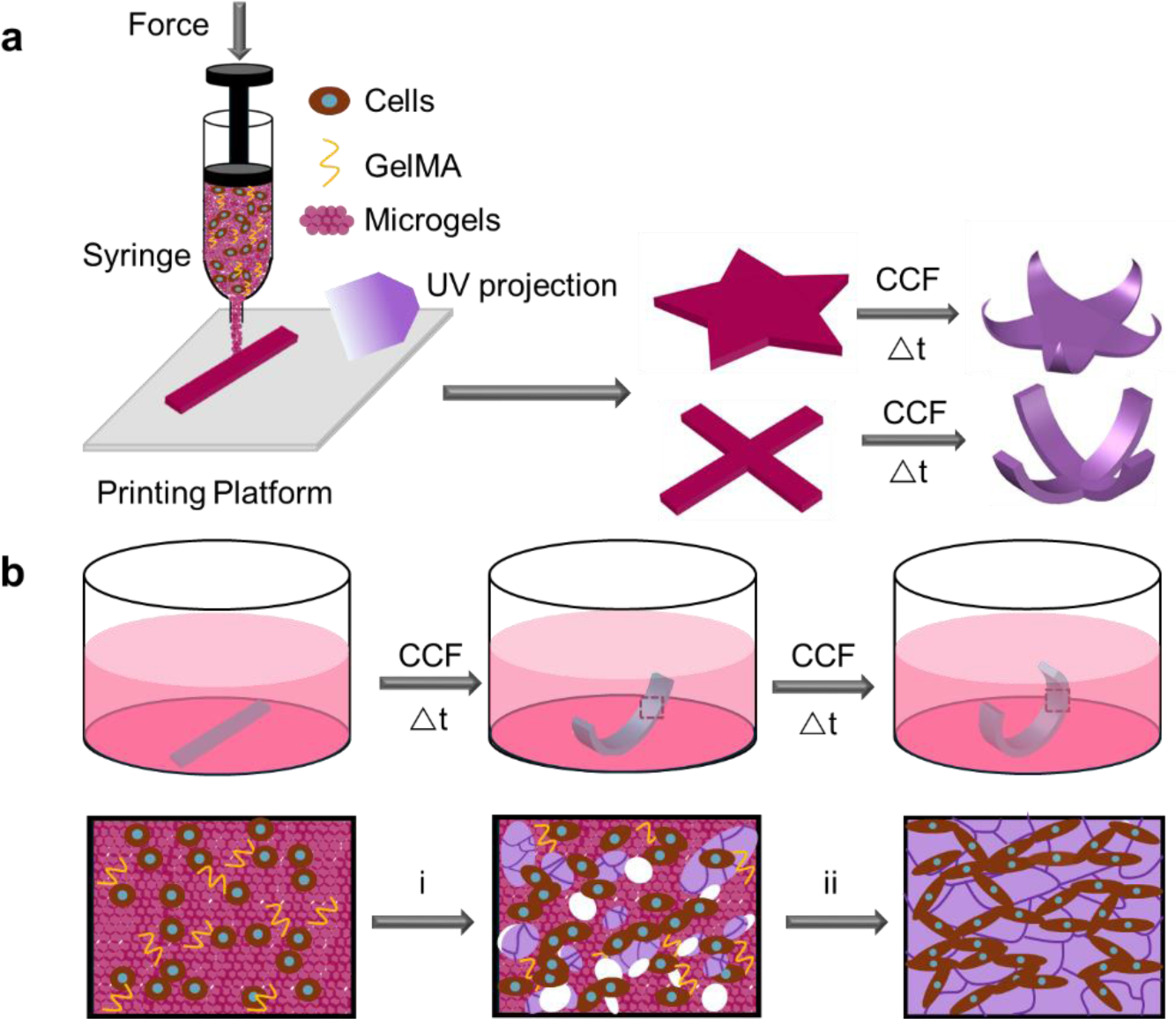
CCF-mediated 4D bioprinting for the creation of geometrically transformable, scaffold-free tissue constructs. (a) Bioprinting of cell-laden bioinks composed of living cells, GelMA, and OMA microgels, followed by CCF-driven shape transformation. (b) Dynamic shape morphing and tissue maturation of living constructs during culture. (i) Microgel degradation and tissue growth; (ii) Complete microgel degradation leading to the formation of tissue-only (scaffold-free) constructs.

## Results and discussion

### Formulation of hydrogel bioinks and rheological characterization

To enable effective shape changes in the bioconstructs, the supporting hydrogel must be sufficiently flexible and mechanically weak to accommodate deformation driven by CCFs. A polymer composition that degrades rapidly after printing is essential to achieve tissue-only 4D condensation while preserving structural integrity during the initial 3D printing process. Therefore, the design of an optimal hydrogel composition is critical.

OMA (Figure S1) was synthesized as the base polymer for cell-laden bioink formulation due to its photocrosslinkablity, high cytocomapbility, and tunable degradability^38,39^. Microgels derived from OMA polymers (OMA microgels) were synthesized with rheological properties tailored for smooth extrusion printing^40,41^. To enhance cell adhesion and interaction with the hydrogel matrix, a small amount of GelMA (3% w/v) (Figure S2) was incorporated into the microgels. GelMA contains bioactive motifs, such as arginine-glycine-aspartic acid (RGD), which promote cell adhesion^42^. After the printing of the composite bioinks, applying photoirradiation could stabilize the printed structures to form robust cell-laden constructs suitable for transfer to and culture in media. It was hypothesized that, following rapid degradation of the OMA matrix, the scaffold would weaken, allowing cells within the construct to exert CCFs and drive shape morphing. This shape change would be facilitated by both the proliferation of cells and the resulting increase in CCFs. Consequently, the composite bioink formulation consisting of OMA microgels and GelMA (OMAGM) was developed for use in this study.

To successfully print freestanding 3D constructs, the composite bioink must exhibit solid-like behavior for structural stability while also possessing shear-thinning properties to ensure smooth extrusion. To evaluate these characteristics, rheological tests were performed on the bioink. A frequency sweep test at 0.1% strain revealed a significantly higher storage modulus (G’) than loss modulus (G’’), indicating a predominantly solid-like structure at low frequencies (0.1–10 Hz) (Figure 1a). To further assess the shear-thinning properties, strain sweep and shear rate ramp tests were conducted. The results showed that G’ remained greater than G’’ across most of the applied strain range, confirming that the bioink retained its structural integrity at low strains (Figure 1b). A crossover between G’ and G’’ at 20% strain marked the transition from a solid-like to a liquid-like state, demonstrating the material’s ability to flow under higher strain conditions. Further analysis of the phase transition was evident from the complex viscosity-shear strain and viscosity-shear rate graphs. The complex viscosity-shear strain curve showed a gradual decrease in viscosity beginning at 4% strain (Figure 1c), while the viscosity-shear rate curve demonstrated a consistent reduction in viscosity with increasing shear rate (Figure 1d). By applying a Power-Law fit to the log-log plot of viscosity versus shear rate within the linear range^43^ from 0.22 s⁻¹ to 2.07 s⁻¹, we determined the flow index (*n*) and the consistency index (*K*) to be 0.26 and 21.66, respectively. These results confirm the bioink’s shear-thinning behavior, which is critical for ensuring sufficient flowability during extrusion.

**Figure 1.**
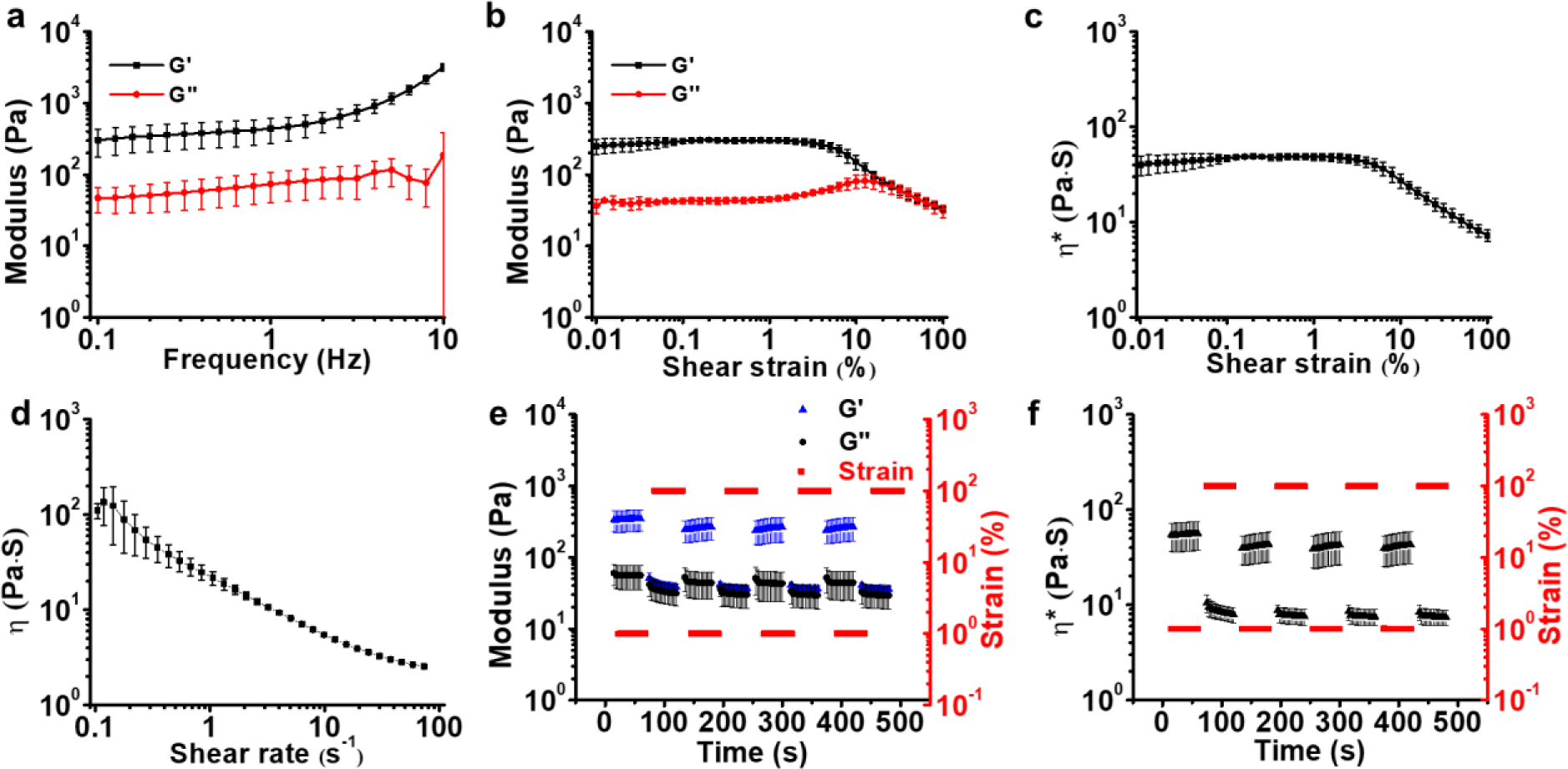
Rheological properties of the OMAGA bioink. (a) Storage modulus (G’) and loss modulus (G’’) as a function of frequency. (b) Changes in G’ and G’’ with increasing shear strain. (c) Complex viscosity (η*) as a function of shear strain. (d) Viscosity (η) as a function of shear rate. (e) Modulus and (f) complex viscosity changes over time under cyclic shear strain of 1% and 100%.

Self-healing is another key parameter for maintaining the stability of printed constructs. After printing, the OMAGM bioink must quickly recover its initial structural state to maintain a stable freestanding form. To evaluate the self-healing ability, a cyclic shear strain test alternating between 1% and 100% strain was conducted. The high strain simulated the shear forces encountered during extrusion, while the low strain represented the post-extrusion state. As shown in Figures 1e and 1f, the bioink exhibited a rapid decrease in modulus and complex viscosity under high strain, followed by an immediate recovery when the strain was released. Minimal hysteresis between cycles indicated that the OMAGM bioinks quickly regained mechanical stability^44^. Even after four cycles, the OMAGM bioink demonstrated consistent recovery capability. Similar rheological performance was observed with the OMA-only bioinks (Figure S3). These results confirm the suitability of both OMAGM and OMA-only bioinks for extrusion-based printing.

### Mechanical characteristics and degradation of 3D printed hydrogel constructs

After confirming the favorable rheological properties for 3D printing, we proceeded with printing using the parameters detailed in Table S1. A hydrogel slab measuring 20 × 20 × 1.0 mm (L × W × H) was initially printed, photocrosslinked (15 s, 6.7 mW/cm²), and then punched into multiple hydrogel discs (d = 8 mm) for testing (Figure 2a). These samples were subjected to dynamic mechanical analysis using frequency sweep mode. The rheogram in Figure 2b shows that both as-prepared OMA and OMAGM hydrogels exhibited a frequency-independent storage modulus (G’), which is a typical characteristic of chemically crosslinked hydrogels^45^. The OMAGM hydrogel had a lower G’ than the OMA hydrogel, suggesting that incorporating GelMA into the OMA matrix resulted in a softer hydrogel.

**Figure 2.**
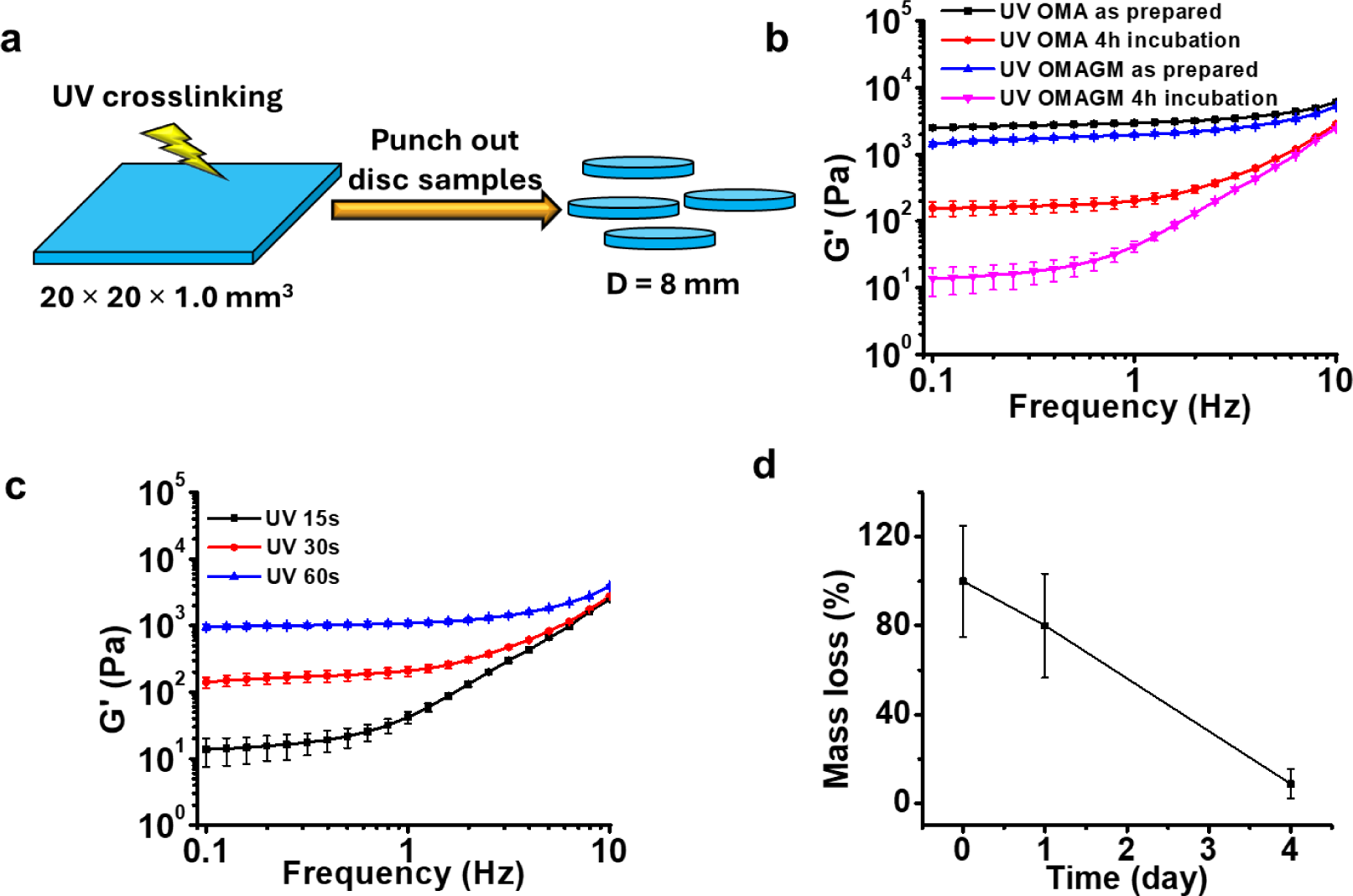
Mechanical properties and degradation behavior of hydrogel constructs. (a) Schematic illustration of hydrogel disc preparation. (b) Storage modulus (G’) of as-prepared and swollen OMA and OMAGM hydrogels as a function of frequency. (c) Storage modulus (G’) of swollen OMAGM hydrogels after varying UV crosslinking time. (d) Mass loss profile of OMAGM hydrogels with a UV crosslinking time of 15 s.

After incubation in cell-expansion media at 37 °C for 4 hours, both OMAGM and OMA-only hydrogels reached a swollen state, which resulted in a significant decrease in G’ and a shift to strong frequency dependence due to networks weakening. The swollen OMAGM hydrogel exhibited a much lower G’ compared to the swollen OMA hydrogel, indicating a weaker matrix due to GelMA incorporation. This reduction in hydrogel stiffness is advantageous for facilitating CCFs to induce deformation. While the as-prepared hydrogels possessed sufficient mechanical rigidity for handling and transferring into culturing media, the marked decrease in stiffness upon swelling is particularly favorable for cell-mediated contraction. Notably, the G’ of the swollen OMAGM hydrogel decreased to below 200 Pa, a value lower than the elastic modulus of most soft tissues^46,47^, thereby reducing mechanical resistance and enabling CCFs to more effectively contract the hydrogel matrix.

The effect of UV crosslinking time on the G’ of the swollen OMAGM hydrogels were further investigated. Increasing the UV crosslinking time enhanced chemical crosslinking, substantially increasing hydrogel stiffness (Figure 2c). A UV crosslinking time of 15 s yielded a weak yet structurally sound hydrogel matrix, with a G’ below 20 Pa, which was chosen for stabilizing the printed construct for further studies. Degradation tests demonstrated that under these conditions, the OMAGM hydrogel rapidly degraded within four days of culture in media (Figure 2d). This rapid degradation not only weakens the hydrogel matrix but also creates space for enhanced cell-cell interactions. Previous study has shown that such interactions can significantly increase cell-traction forces (or CCFs)^48^. Therefore, rapid degradation may enhance the ability of CCFs to contract the hydrogel matrix, acting as a key mechanism for construct deformation and the formation of scaffold-free structures.

### Examination of the roles of live cells in driving 4D shape transformation

In soft hydrogels like collagen, live cells have been shown to induce effective matrix contraction through CCFs, with fibroblasts identified as a robust cell type for these studies^49,50^. In this study, the role of live cells was explored in driving 4D shape transformation via controlled matrix contraction, using NIH3T3 fibroblasts as a model. cell-laden hydrogel constructs using OMA and OMAGA bioinks were printed. The experimental setup included three groups of NIH3T3-laden hydrogels, each with a cell density of 1 × 10^8^ (100M) cells/mL bioink: OMA, OMAGM, and an OMAGM loaded with dead cells serving as a control. Disc-shaped constructs, prepared as illustrated in Figure 2a, were cultured in cell-growth media for 14 days. Live/dead staining conducted 4 hours post-fabrication indicated high cell viability within OMA and OMAGM with live cells constructs, whereas cells in the dead control constructs were completely nonviable (Figure S4). Visually, the OMA and OMAGM disc constructs with live cells progressively curled upward to form bowl-shaped structures (Figure 3a). In contrast, the dead control constructs maintained a flat shape throughout the culture period until they eventually collapsed due to hydrogel degradation. The OMA and OMAGM constructs with live cells remained structurally integrated over the 14 days, forming well-morphed constructs (Figure 3b). Live/dead staining revealed high cell viability, a marked increase in cell density, and the formation of dense cellular networks in these constructs as the culture period extended from D0 to D14 (Figure 3c). H&E staining over 14 days of culture revealed that cells within the OMAGM constructs progressively formed into well-organized structures (Figure 3d and S5), with no residual hydrogel visualized at D14, indicating that only cells and ECM remained. Interestingly, the cells at the constructs’ rim appeared more densely organized, potentially contributing to differential contraction along the radial direction and, consequently, the observed curling (Figure 3e).

**Figure 3.**
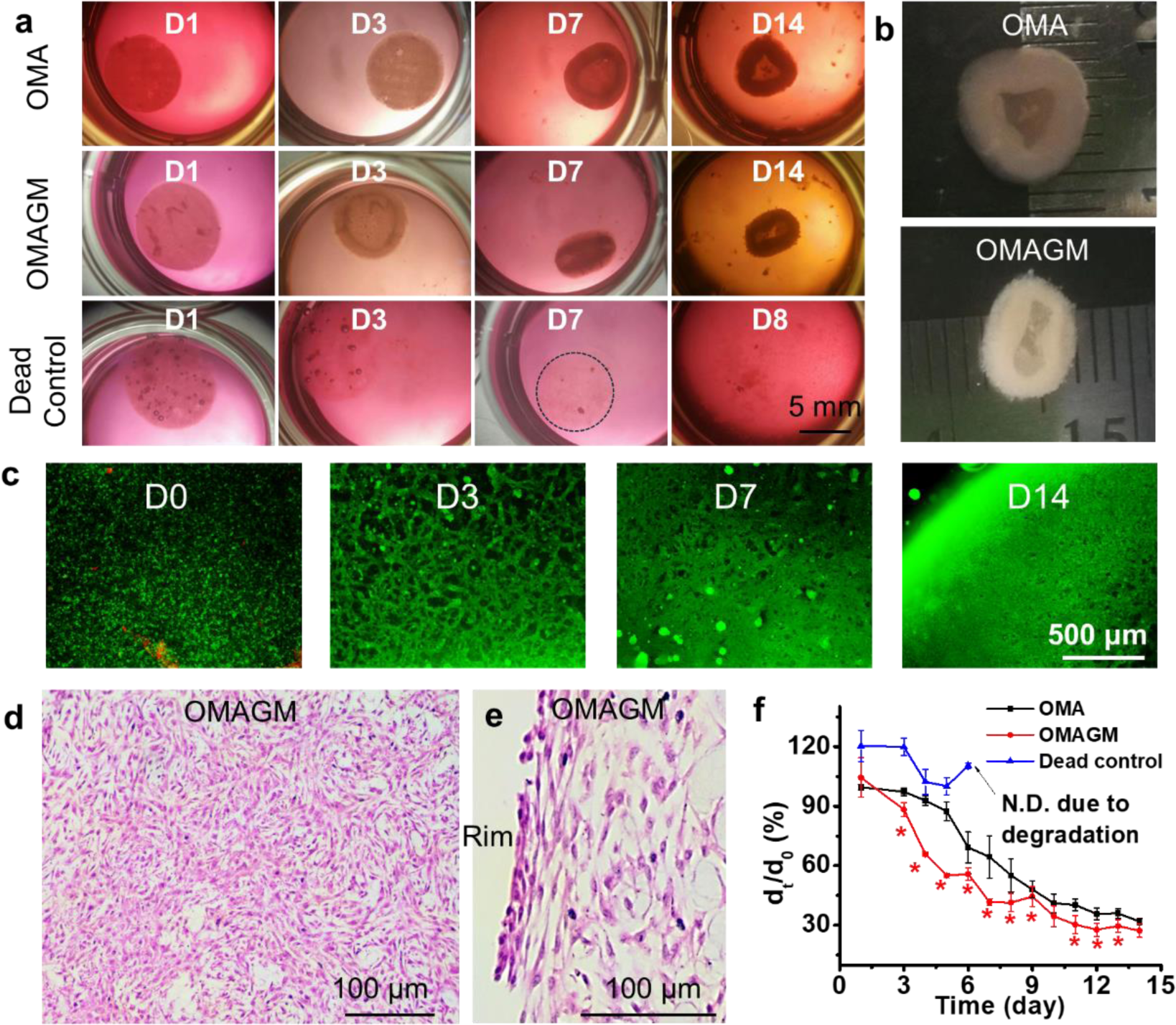
Roles of live cells in driving 4D shape transformation. (a) Time-lapse images of various constructs on days 1, 3, 7, and 14 of culture. Note that the dead cell constructs disintegrated completely by D7 and are not shown thereafter, with only loose polymer fragments remaining in the dish. (b) Morphed tissue-only constructs derived from the OMA and OMAGM groups. (c) Live/dead staining images of cell-laden OMAGM constructs at different times. (d and e) H&E staining images of the tissue-only construct formed from the OMAGM group. (f) Change in diameter ratio (d_t_/d_0_) over culture time, where d_t_ is the diameter of the disc construct on a given day, and d_0_ represents the initial diameter on D0 (8 mm). *p < 0.05 compared to groups of OMA and dead control. Construct fabrication parameters - cell density: 100M; UV: 15s at 20 mW/cm^2^; disc dimensions: d_0_ = 8.0 mm, h = 1.0 mm.

We then quantified the deformation and outlined the deformation profiles of the three groups, following the protocols described in Figure S6, with results shown in Figure 3f. The quantified deformation data aligned with visual observations, showing more rapid and pronounced contraction in OMAGM constructs compared to OMA. Previous studies on cell-laden collagen hydrogels showed the primary mechanism driving hydrogel contraction is the CCFs exerted on the matrix during cell attachment and locomotion^51,52^. The presence of GelMA in OMAGM likely facilitates stronger initial cell adhesion the hydrogel matrix, enabling better transfer of cell contraction forces and more significant and rapid shape deformation than in OMA constructs. Conversely, in the dead control samples (dead cell-laden OMAGM), the inability of dead cells to adhere to the hydrogel or secrete ECM led to rapid construct degradation and disintegration within a week. Interestingly, the degradation of these dead cell samples was slower than that of empty OMAGM constructs (Figure 2d), possibly due to the large cell quantity within them.

Previous studies on cell-induced contraction in soft collagen hydrogels have shown homogeneous matrix contraction, likely due to the large pore sizes in collagen hydrogels, which allow easy reorganization by cell traction and locomotion^53-56^. Consequently, only in-plane shrinkage of construct shapes was observed in those studies. In this study, the mechanically sound yet degradable OMAGM matrix provided temporary mechanical resistance, inducing anisotropic contraction during the initial days of culture. This facilitated observable deformation, underscoring the crucial role of a structurally stable but degradable hydrogel matrix in designing CCF-inducible, shape-transformable hydrogel constructs. However, the specific mechanisms underlying this anisotropic contraction remain unclear at this stage. One possible explanation is that hydrogel regions closer to the edge exhibited a faster degradation profile due to greater accessibility to the culture media, leading to enhanced media exchange and accelerated hydrolysis. Consequently, cells in these regions generated stronger collective contractile forces, inducing anisotropic contraction along radial directions. Along with the restriction imposed by the well plate at the bottom, this contraction resulted in the hydrogel disc curling upward into a bowl-like shape. While GelMA accelerates shape deformation, OMA, despite lacking active cell-adhesive bio-ligands, undergoes rapid degradation, generating pores that weaken the mechanical integrity of the matrix and facilitate cell-cell interactions. Consequently, effective shape deformation was also observed in cell-laden OMA-only constructs.

### Impact of cell density, hydrogel stiffness, and construct dimensions on shape-morphing capability

Having established the critical role of CCFs in driving the shape transformation of these cell-laden constructs, we systematically investigated the effects of cell density, hydrogel stiffness, and construct dimensions on shape-morphing. Initially, we screened cell densities ranging from 20 to 200M cells/mL of bioink. Constructs were subjected to live/dead staining after 3 hours of culture to assess cell viability. Cells within the constructs remained highly viable across all cell densities (Figure S7). Constructs with starting cell densities of 20, 50, and 100M cells/mL displayed increasing cell retention, while those with 200M cells/mL exhibited a significant reduction in cell retention. This reduction is likely due to the inability of the polymer network to sufficiently entrap cells at such a high density, resulting in the leakage of cells into the culture media within the first few hours. Consequently, constructs from the 20, 50, and 100M cells/mL groups were further cultured for three days. As observed in Figure S8, constructs with 20M cells/mL collapsed by D3, unable to maintain their integrity due to insufficient cell condensate formation following rapid hydrogel degradation, similar to the dead control samples. In contrast, constructs with 50 and 100M cells/mL retained their structure, with the 50M cells/mL constructs exhibiting greater deformation. However, the constructs with a cell density of 50M cells/mL were too fragile to be moved out of the plate well, prompting us to determine that a density of 100M cells/mL is optimal for subsequent studies.

To investigate the impact of hydrogel stiffness on construct morphing, constructs were prepared with varying photocrosslinking times, ranging from 15 to 60 s. Constructs subjected to longer photocrosslinking times exhibited increased rigidity. Consequently, cell-laden constructs with 30 s and 60 s photocrosslinking times showed minimal contraction and limited upward curving during the two-week culture period (Figures 4a,b). Interestingly, during the initial five days, constructs with a 15 s photocrosslinking time exhibited less contraction compared to those with 30 s and 60 s times. This difference is attributed to the higher initial swelling of the 15 s constructs, resulting in greater volume expansion. After D5, CCFs induced significant contraction in the 15 s constructs, leading to a smaller construct size and more pronounced morphing compared to the 30 s and 60 s constructs.

**Figure 4.**
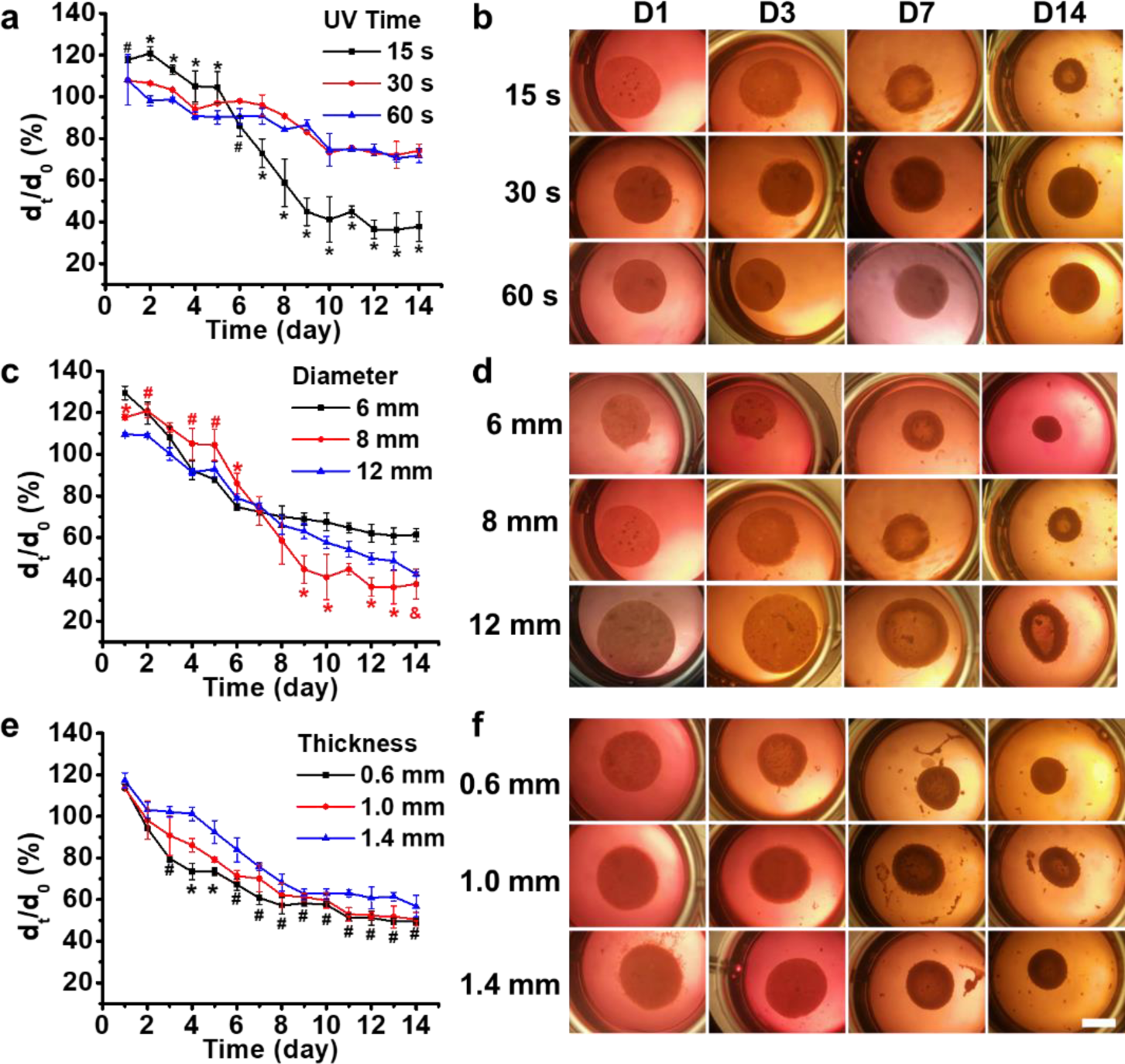
Impact of hydrogel stiffness and construct dimensions on morphing capacity. Effect of (a,b) hydrogel stiffness (photocrosslinking time), (c,d) construct diameter, and (e,f) construct thickness on construct morphing. For (a): *p < 0.05 compared to 30 s and 60 s constructs, ^#^p < 0.05 compared to 30 s construct. Scale bar: 5 mm. For (c): *p < 0.05 compared to 6 mm and 12 mm constructs, ^#^p < 0.05 compared to 12 mm construct, ^&^p < 0.05 compared to 6 mm construct. For (e): *p < 0.05 compared to 1.0 mm and 1.4 mm constructs, ^#^p < 0.05 compared to 1.4 mm construct. Unless otherwise specified, the preparation parameters for all constructs were as follows: cell density: 100M cells/mL bioink; UV exposure: 15 s at 20 mW/cm²; disc dimensions: d_0_ = 8.0 mm, h = 1.0 mm.

Next, we examined the impact of construct dimensions, including diameter and thickness, on shape morphing (Figure 4b-e). All constructs underwent rapid contraction and shape morphing over 14 days of culture. However, among constructs of varying diameters, the 8 mm constructs exhibited greater contraction after D8 than the 6 mm and 12 mm constructs and maintained their bowl-like shape by D14. In contrast, the 6 mm constructs formed cell clusters without a defined shape by D14 (Figure 4b,c). Rapid contraction and shape morphing across constructs with different thicknesses was also observed (Figure 4c,e). Among these, constructs with the smallest thickness of 0.6 mm displayed the most pronounced and rapid contraction and morphing. However, by D14, these constructs lost their shape, forming cell clusters. In comparison, 1.0 mm and 1.2 mm constructs maintained well-defined bowl-like shapes by D14.

The findings indicate that construct morphing is governed by a complex interplay of factors, including cell density, hydrogel stiffness, and construct dimensions. These parameters collectively offer a multifactorial strategy for programming the morphing capabilities of cell-laden constructs. When designing CCF-4D constructs, it is essential to strike a balance between the desired morphing potential and the ability to preserve a well-defined shape over extended culture periods. This dual consideration is critical for achieving functional and stable tissue constructs in advanced 4D tissue engineering applications.

### Impact of initial geometry on the shape-morphing capacity

Given the demonstrated capability of CCF-mediated shape morphing in disc-shaped constructs, whether CCF could effectively drive shape changes to form different complex architectures was investigated by varying the initial geometric designs (Figure 4). Consistent with the findings from disc-shaped constructs, these constructs exhibited continuous and dynamic shape morphing throughout the culture period. Specifically, strip-shaped constructs bent into a “C” pattern, reaching peak curvature at D7, followed by partial relaxation by D14 (Figure 4a). Similarly, sheet constructs rolled into tubular forms (Figure 4b), while the 4-arm gripper geometry curled inward, resulting in a flower-like configuration (Figure 4c). A comparable morphing process was observed in star-shaped constructs (Figure 4d). These results underscore the potential of leveraging initial geometric design to generate complex tissues through CCF-driven shape morphing. This approach highlights the effectiveness of CCF in directing the programmed shape morphing of cell-laden constructs, paving the way for biomimetic 4D engineering that replicates the dynamic morphogenesis observed in tissue development.

**Figure 4.**
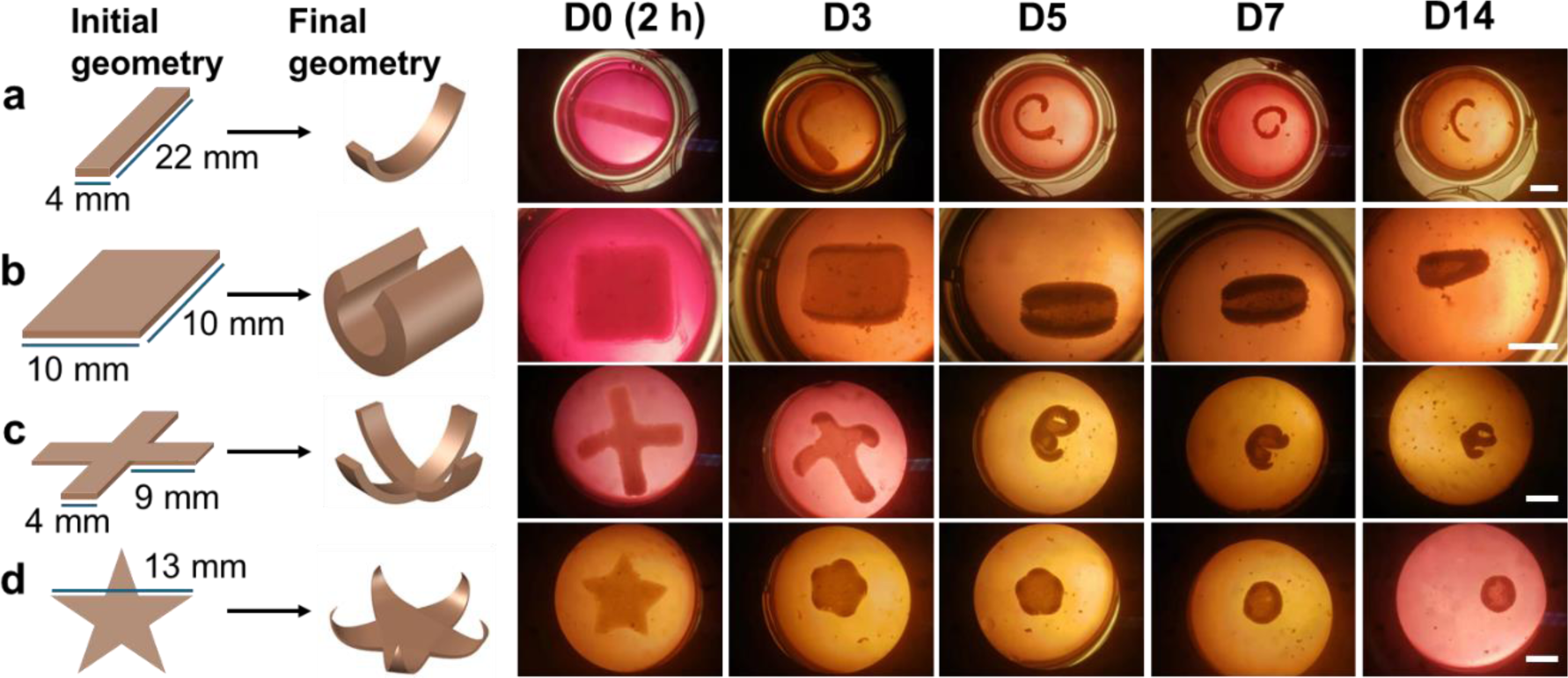
Shape morphing of cell-laden constructs with varying initial geometries over 14 days of culture in media.: (a) strip-shaped, (b) square sheet-shaped, (c) four-arm gripper-shaped, and (d) star-shaped. Construct fabrication parameters: cell density: 100M cells/mL bioink; UV exposure: 15 s at 20 mW/cm². Construct thicknesses: 0.6 mm. Scale bars: 5 mm.

### 4D tissue engineering

In the human body, tissue development is inherently tied to morphological evolution, characterized by the formation of specific architectures through a dynamic morphing process that includes buckling, folding, elongation, curling, and/or wrinkling. This morphogenic process is crucial in the development of complex tissue curvatures observed in structures such as intestinal villi and epithelial tubes^57^. Not only does this process expand structural complexity to facilitate functionality, such as increased surface to volume ratio, as seen in branching morphogenesis, but it also provides biomechanical cues that are vital for guiding tissue development^58,59^. Therefore, introducing dynamic elements that mimic the morphological dynamics of native tissues may be valuable for advancing tissue engineering practices to a horizon of more effective tissue formation and to gain a deeper understanding of developmental processes.

To evaluate the ability of CCFs to drive shape morphing in engineering tissues, multipotent human mesenchymal stem cells (hMSCs) within printed cell-laden constructs were cultured in tissue-specific environments to promote tissue maturation. Initially, hMSC chondrogenesis were induced within chondrogenic differentiation media to engineer cartilage-like tissues using a simple strip-shaped prototype (termed Exp group). A control group (Ctrl) was maintained in normal growth medium. Over the culture period, both Ctrl and Exp constructs exhibited continuous morphological evolution, with the Exp constructs demonstrating more rapid and pronounced volume shrinkage compared to the Ctrl group (Figure 5a,b). However, no statistical significances were observed in the bending angle (Figure 5c). By D10, the Exp constructs had transformed from a “C” shape into a “kidney bean” shape, while the Ctrl constructs maintained their “C” geometry. The more pronounced geometrical changes in the Exp group may be attributed to the differentiated chondrocytes exerting stronger CCFs compared to the undifferentiated hMSCs in the Ctrl group^60,61^. Biochemical assays revealed significantly higher glycosaminoglycan (GAG) production normalized to DNA content (GAG/DNA) in Exp constructs compared to Ctrl constructs (Figure 5d,S9), indicative of robust chondrogenesis. These findings were corroborated by histological analyses using Safranin O (SafO) and Toluidine Blue O (TBO) staining (Figure 5e and f).

**Figure 5.**
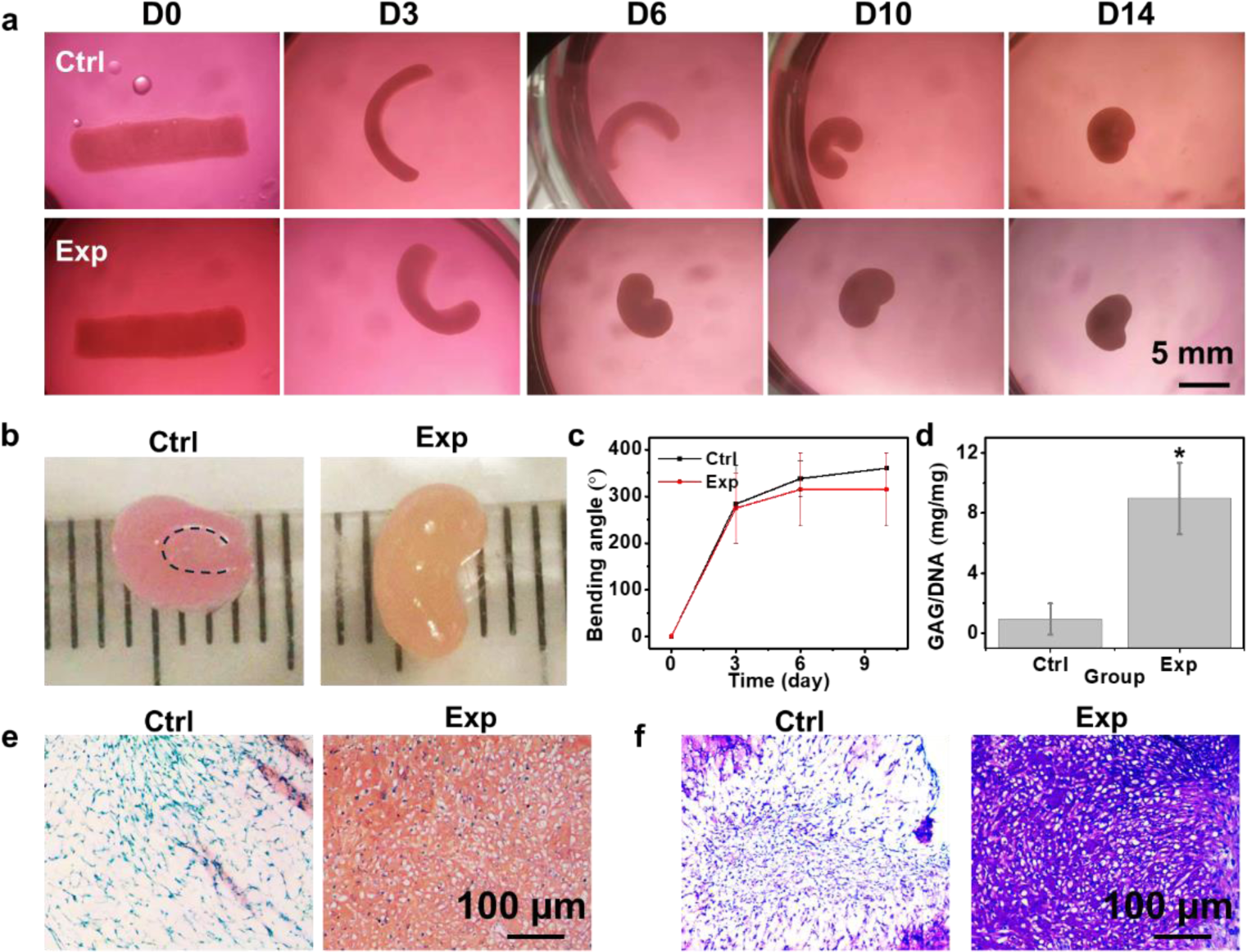
Differentiation study of 4D constructs for chondrogenesis. (a) Images of constructs at different times during a 14-day culture period. (b) Images of D14 constructs removed from culture media, with each ruler tick mark representing 1 mm. The dotted line in the Ctrl sample outlines the curvature arc. (c) Quantitative analysis of the bending angle change in Ctrl and Exp groups over time. (d) Biochemical analysis of GAG production normalized to DNA content (GAG/DNA) in Ctrl and Exp constructs at D14. *p < 0.05. Histological staining of D14 constructs using (e) SafO and (f) TBO. Ctrl represents the control group cultured in growth medium, while Exp denotes the group cultured in chondrogenic differentiation medium.

Osteogenic differentiation studies further validated the CCF-4D engineering of bone tissues. Constructs cultured in media displayed progressive shape changes over a 21-day period (Figure 6a). They formed forming distinct “C” shapes by D6. However, by D14, the Ctrl constructs gradually lost this geometric characteristic by D14, whereas the Exp constructs underwent significant shrinkage, retaining a small “C” shape. The Exp constructs, subjected to osteogenic differentiation, exhibited less pronounced bending angles compared to the Ctrl group throughout the culture period (Figure 6b), likely due to the production of mineralized ECM, which increased tissue rigidity and rapidly balanced the stress between cells and their surrounding matrix. This rigidity led to reduced tissue deformation in the Exp constructs, even though differentiated osteoblasts, known for their larger size, can exert higher traction forces compared to undifferentiated stem cells^62,63^. Biochemical analysis at D21 confirmed the differentiation capabilities within the osteogenic media, as evidenced by significantly elevated levels of alkaline phosphatase (ALP) and calcium (Figure S10), both normalized to DNA content (ALP/DNA and Ca/DNA, respectively), in the Exp group relative to the Ctrl group (Figure 6c and d). Histological assessments using H&E and Alizarin Red S (ARS) staining further demonstrated denser tissue organization and more intense red staining in the Exp constructs, indicative of neo-bone-like tissue formation (Figure 6e and f).

**Figure 6.**
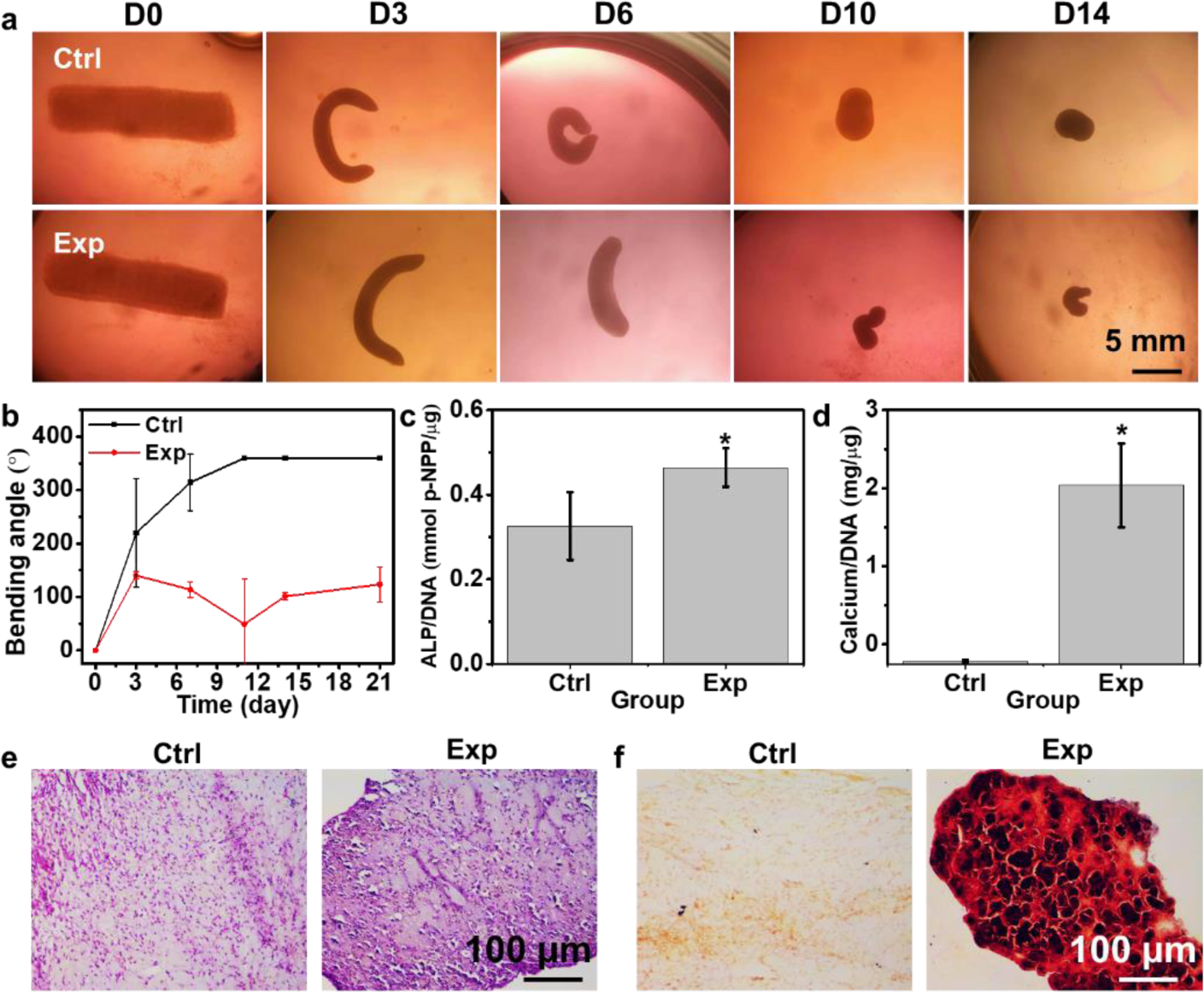
Differentiation study of 4D constructs for osteogenesis. (a) Images of constructs at different times during a 14-day culture period. (b) Quantitative analysis of the change in bending angle for Ctrl and Exp groups over time. Quantification of (c) ALP activity normalized to DNA (ALP/DNA) and (d) calcium content normalized to DNA (Ca/DNA) in Ctrl and Exp constructs at D14. *p < 0.05. Histological staining of D21 constructs using (e) H&E and (f) ARS. Ctrl represents the control group cultured in growth medium, while Exp denotes the group cultured in osteogenic differentiation medium.

Collectively, these chondrogenic and osteogenic studies highlight the ability of the 4D constructs to support differentiation while undergoing geometric transformations, thereby affirming their potential as models for tissue development. Throughout tissue maturation, cells within the transient hydrogel matrix actively remodel the ECM, ultimately forming complex tissues in the absence of hydrogel materials. This dynamic process closely mimics the native tissue development observed in vivo, underscoring the value of coupling 4D morphogenesis with tissue engineering to better replicate the intricacies of *in vivo* tissue formation.

Compared to conventional 4D systems that primarily rely on physical and chemical stimuli to elicit shape morphing, leveraging biological forces, such as CCFs generated by live cells, represents a significant advancement in biomimetic 4D tissue engineering. However, due to the inherently weak nature of CCFs, existing CCF-driven 4D systems often require ultrasoft hydrogel matrices to enable detectable shape transformation^36,37,64^. While these soft hydrogels facilitate cellular force transmission, they pose challenges in forming stable, freestanding structures, necessitating the use of a mechanical support to maintain the integrity of the initial cell-laden constructs. To overcome this limitation, our recent work on CCF-driven 4D systems introduced a mechanically self-adaptive hydrogel scaffold incorporating sacrificial gelatin microspheres^61^. This design allows for the fabrication of stable 3D freestanding structures that undergo controlled softening during culture as the gelatin microspheres dissolve. This progressive softening amplifies the impact of CCFs, enabling significant shape transformations over time. While hydrogel materials play a critical role in maintaining structural integrity during culture and differentiation, their prolonged presence could interfere with tissue remodeling and introduce potential immunotoxicity when applied in vivo. To address these challenges, we herein have developed a mechanically robust yet rapidly degradable hydrogel system that supports stable 3D printing while facilitating the formation of cell condensates capable of programmed 4D shape morphing. Upon culture, this system enables the formation of complex cell condensate architectures devoid of residual hydrogel biomaterials, offering a CCF-based dynamic platform that closely mimics the morphogenesis of native tissue development. However, in this system, the morphing directionality is primarily dictated by the spatial restriction imposed by the plate bottom, lacking a mechanism to precisely control morphing directions, thereby limiting the range of achievable morphologies. Incorporating signaling molecules or spatially aligning cell positions could provide a means to regulate CCF directionality in future studies, enabling more precise and programmable shape transformations to better emulate complex morphogenic processes.

## Conclusions

This work presents an innovative 4D bioprinting platform that uniquely utilizes CCF as the exclusive stimulus to orchestrate shape transformation. The bioink, composed of degradable OMA microgels integrated with cell-adhesive GelMA, was formulated to ensure smooth cell loading and printing. The engineered hydrogel scaffolds served as temporary mechanical supports, maintaining structural integrity while rapidly degrading in synchrony with morphogenesis during culture. This process facilitated the generation of tissue constructs with complex, deformed architectures, devoid of residual polymeric materials. By precisely programming CCF within geometrically defined constructs, global shape transformation in large constructs was achieved, resulting in the formation of tissues with complex architectures. Utilizing this unique CCF-driven 4D bioprinting platform, 4D morphogenesis through proof-of-concept studies in cartilage and bone tissue engineering was demonstrated. This CCF-4D bioprinting approach represents a significant advancement in regenerative medicine, with broad potential applications in biomimetic tissue engineering and beyond.

## Methods

### OMA microgel (OMA MG) synthesis

Oxidized methacrylate alginate (OMA) was synthesized according to previously published protocols^38^. Briefly, 10 g of alginate was dissolved in 900 mL of deionized Milli-Q water overnight. Next, 0.545 g sodium periodate solution in 100 mL of deionized water was added to the alginate solution. The mixture was kept in the dark for 24 hours under stirring at room temperature. Subsequently, 19.52 g of 2-(N-morpholino)ethanesulfonic acid (MES, Sigma, Cat#M8250-1KG) and 17.53 g of sodium chloride were added, and the pH was adjusted to 6.5 using 5N sodium hydroxide (NaOH). Then, 0.589 g of N-hydroxysuccinimide (NHS, VWR, cat#102614-812) and 7.776 g of 1-ethyl-3-(3-dimethylaminopropyl)carbodiimide hydrochloride (EDC·HCl, Oakwood, cat#024810-250g) were then added under vigorous stirring. After 10 minutes, 0.844 g of 2-aminoethylmethacrylamide hydrochloride (AEMA, PolySciences, cat#21002-10) was then added, and the solution was left to react for 24 hours under stirring in the dark. After precipitation using chilled acetone, the alginate was rehydrated and dialyzed against deionized Milli-Q water for the next 3 days using MWCO 3500 Da dialysis tubing (Spectra, cat#08-670-5B). The product was treated with charcoal (Neta Scientifics, cat#099536) for 30 minutes and then lyophilized for 14 days. The structure of OMA was confirmed using ^1^H NMR analysis on the Bruker 600 MHz AVANCE III NMR Spectrometer, and shown in **Figure S2**. The actual methacrylation was determined to be 9.1%.

The OMA polymer (2 g) was dissolved in deionized water (100 mL) at 2% w/v solution and then crosslinked in a 2 M CaCl_2_ solution (1 L) for 12 hours. Next, the OMA hydrogel was collected and fragmented finely in a blender (Osterizer MFG, at “pulse” speed) for 5 minutes in the presence of 70% ethanol (EtOH, 100 mL). The mixture was then centrifuged and stored in 70% EtOH at - 20 °C for future use.

### GelMA synthesis

GelMA was synthesized according to previously published protocols^65,66^. Briefly, 20 g of Gelatin Type B (Sigma, cat#G9391-100G) was dissolved in PBS (pH 7.4). Then 10 mL of methacrylic anhydride (Sigma, cat#276685-500ML) was added to the solution slowly (1 mL/min) while stirring vigorously. The reaction was maintained for 1 hour at 50 °C with stirring, and then at room temperature overnight at a pH of 9. The next day, GelMA was precipitated in excess acetone at 4 °C, and then purified by dialysis (MWCO 12-14k Da, Spectrum Laboratories Inc., cat#132709) against deionized Milli-Q water for 10 days at 50 °C. The final product was obtained by lyophilization for 10 days.

### Cell culture

NIH 3T3 fibroblasts (ATCC) were expanded in growth media composed of Dulbecco’s Modified Eagle Medium-low glucose (DMEM-LG, Sigma, cat#D5523-50L) with 10% fetal bovine serum (FBS, Sigma, cat#F19) and 1% penicillin/streptomycin (P/S, Gibco, cat#15140122) until the cells reached 85% confluency. Media was changed every other day. On the day of printing, the cells were harvested and suspended at 1 × 10^8^ cells/mL composite bioink. After printing, the constructs were cultured in Dulbecco’s Modified Eagle Medium-high glucose (DMEM-HG, Sigma, cat#D7777-10x1L) supplemented with 10% FBS and 1% P/S.

### Cell laden composite bioink preparation

On the day of printing, the OMA MGs stored in 70% EtOH was washed four times with deionized water containing 0.05% w/w photointiator (2-hydroxy-4’-(2-hydroxyethoxy)-2-methylpropiophenone, Sigma, cat# 410896-10g), followed by a wash with DMEM-HG containing 0.05% w/w photoinitiator. Lyophilized GelMA was then added to the recovered OMA MGs at a concentration of 3% w/v and vortexed vigorously to yield OMAGM bioinks. Next, cells added to the OMAGM at a concentration of 1 × 10^8^ cells/mL bioink. The cell-laden bioink in a syringe was mounted on a BIOX 3D printer (Cellink, San Diego, CA) and was then printed using the parameters listed in Table S1 in the Supporting Information file. After printing, the constructs were further UV-crosslinked (20 mW/cm^2^) for a specific duration. The constructs were then transferred into culture media and placed in an incubator for further investigation. Disc-shaped constructs were cultured in 24-well plates with 2 mL of media, while strip, square, gripper, and star-shaped constructs were cultured in 12-well plates with 4 mL of media. Differentiation strip constructs were cultured in 6-well plates with 8 mL of media. In all cultures, half of the media volume was changed daily.

### 4D printed chondrogenesis constructs

In this study, hMSCs were isolated and harvested according to previously established protocol[67]. hMSCs at passage 4 were used to mix with OMAGM to generate hMSCs-laden bioinks (1 × 10^8^ cells/mL bioink), which were then printed into constructs and crosslinked as described above. The constructs were then cultured in chondrogenic medium composed of DMEM-HG (Sigma, cat#D5648-10X1L), 1% v/v P/S, 10% v/v ITS^+^ (BD Biosciences, cat#354352), 1% v/v nonessential amino acids (NEAA, Gibco cat#11140050), 100 mM sodium pyruvate (Fisher, cat#SH3023901), 10^-7^ M dexamethasone (Sigma, cat#D4902-100mg), 129 nM L-ascorbic acid phosphate (Wako USA, cat#013-12061), and 10 ng/mL transforming growth factor beta-1 (TGF-β1, Peprotech, cat#100-21-10UG). The culture medium was changed every day. A control group was cultured in normal growth medium (DMEM-HG supplemented with 10% FBS and 1% P/S) for comparison. The constructs were cultured for 14 days until collection. Sample sizes of *N = 4* were collected for biochemical analysis and *N = 2* were collected for histological staining.

### 4D printed osteogenesis constructs

The hMSCs at passage 3 (1 × 10^8^ cells/mL bioink) were used to prepare the cell-laden OMAGM bioinks for printing. The printed constructs were cultured in differentiation media composed of DMEM-HG, 100 ng/mL bone morphogenic protein-2 (BMP-2, Genscript, cat#Z02913), 10% v/v FBS, 1% v/v P/S, 100 µM dexamethasone, and 100 µM L-ascorbic acid phosphate. Constructs in the control group were cultured in normal growth media for comparison. The constructs were cultured for 14 days until collection. Sample sizes of *N = 4* were collected for biochemical analysis and *N =2* were collected for histological staining.

### Histology

Staining was performed according to previously established protocols^67,68^. Briefly, samples were submerged in 10% neutral buffered formalin at 4 °C overnight. The samples were then dehydrated, paraffin-embedded, and sectioned at 7 µm thickness using a Leica RM2255 microtome (Leica Biosystems, Nussloch, Germany). Next, the samples were stained with Mayer S Hematoxylin (Fisher, cat#TA125MH) and Eosin (Electron Microscopy services, cat#26763-03) (H&E) to observe cell morphology and tissue organization. To examine GAG production, a combination of 0.1% w/v Safranin O (SaFO, Acros Organics, cat#477-73-6) and counterstain of 1% w/v Fast Green (Fisher, cat#F99-10), or Toluidine Blue (TBO, Fisher, cat#92-31-9) staining were used^67^. Briefly, SafO was applied for 5 minutes followed by 2 minutes of Fast Green. TBO, at pH 0f 4.0, was applied for 30 minutes on the samples. Additionally, Alizarin Red S (Sigma, cat# A5533-25G) staining was used to examine calcium mineralization deposits according to a previously reported protocol^69,70^.

### Live/dead cell staining

Live/dead cell staining was performed at predetermined timepoints during culture for assessing cell viability using a previously established protocol^71^. Briefly, constructs were stained with 16 µL of fluorescein diacetate (2 mg/mL, Sigma, cat#F7378) and 8 µL of propidium iodide (2 mg/mL, Sigma, cat#P4170-25MG), respectively for 3 minutes, and then imaged on a Nikon Eclipse TE300 fluorescence microscope (Nikon, Tokyo, Japan) equipped with a AmScope MU1403 camera (AmScope, Irvine, California). *N = 3*.

### Biochemical assays

DNA, GAG, ALP, and calcium contents were quantified according to the modified protocols from previous literature^68^. Briefly, prior to quantification, chondrogenesis samples were digested in 0.5 mL of a solution containing 50 µg/mL papain (papaya latex, Sigma, cat#3P4762), 2 mM L-cysteine (Sigma, cat# C7352), 50 mM sodium phosphate (Sigma, cat#BP332-500), and 2 mM EDTA (Fisher, cat# BP120-1, pH 6.5) for 12 hours at 65 °C. Next, 0.5 mL of nuclease-free water was added to each sample solution, and the solution was vortexed for 1 minute. DNA concentration was determined using a PicoGreen (Invitrogen, cat# P7581) dsDNA fluorescence assay on a plate reader (Molecular Devices ID5) with excitation at 480 nm and emission at 520 nm. GAG content was determined using a 1,9-dimethylmethylene blue (DMMB) (Sigma, cat# 341088) absorbance assay at 595 nm on the plate reader. *N = 4*.

Prior to quantification, osteogenesis samples were homogenized in 1 mL of Cellytic Buffer M (Sigma, cat# C2978-250ML) for 90 s on ice. Next, the homogenates were centrifuged at 500× g for 5 minutes. DNA content was determined using the same assay as described for chondrogenesis samples. ALP content was quantified using an absorbance assay at 405 nm on the plate reader. Calcium content was determined using a calcium assay solution kit (Pointe Scientific, cat#C7503-480) with absorbance measured at 570 nm on the plate reader. *N = 4*.

### Bending angle calculations

The bending angle was quantified according to a previously published protocol^23^. Briefly, the hydrogel bar was extended into a circle with equal crosshatches that divide the circle into four quadrants. The hydrogel bar with the circle and crosshatches was then loaded into NIH ImageJ to determine the angle measurement. If the hydrogel bar curved past the half of the circle, the resulting bending angle was subtracted from 360° to get the bending angle. *N = 3*.

### Statistical analysis

All graphs are reported as mean ± standard deviation (±SD). Statistical analsysi was performed and significance was determined using one-way ANOVA with a post-hoc Tukey HSD test. p < 0.05 was considered significant unless otherwise specified.

## Acknowledgments

The authors gratefully acknowledge funding support from the Department of Veterans Affairs, Veterans Health Administration, Office of Research and Development, Rehabilitation Research and Development Service under award numbers RX004288 and RX004825 and the National Institutes of Health’s National Institute of Arthritis and Musculoskeletal and Skin Diseases under award number R01AR081448. The contents of this publication are solely the responsibility of the authors and do not necessarily represent the official views of the Department of Veterans Affairs or the National Institutes of Health. The authors also acknowledge Hudson Gasvoda’s guidance on using AUTOCAD 2024 for drawing schematics.

## CRediT authorship contribution statement

**A.D.**: Conceptualization, Methodology, Investigation, Formal Analysis, Visualization, Writing-Original Draft, Writing – Review & Editing. **K.L.G.**: Methodology, Investigation, Formal Analysis, Visualization, Writing-Original Draft, Writing – Review & Editing. **D.S.C.**: Methodology, Investigation, Formal Analysis, Visualization. **E.A.**: Conceptualization, Methodology, Formal Analysis, Resources, Supervision, Funding acquisition, Writing – Review & Editing.

