## Supporting Information for "Cell Contractile Force-Mediated Morphodynamical Tissue Engineering via 4D Printed Degradable Hydrogel Scaffolds"

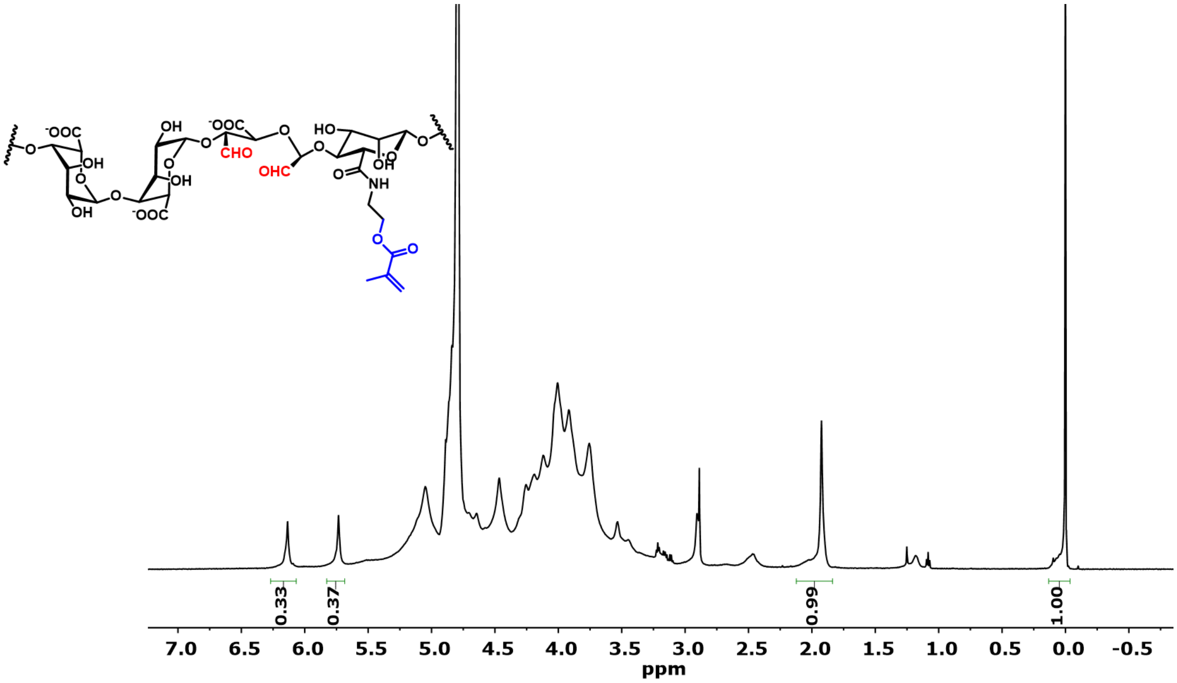


**Figure S1**. Molecular structure and $^1$H NMR ($D_2O$, 500 MHz) of O5M20A.

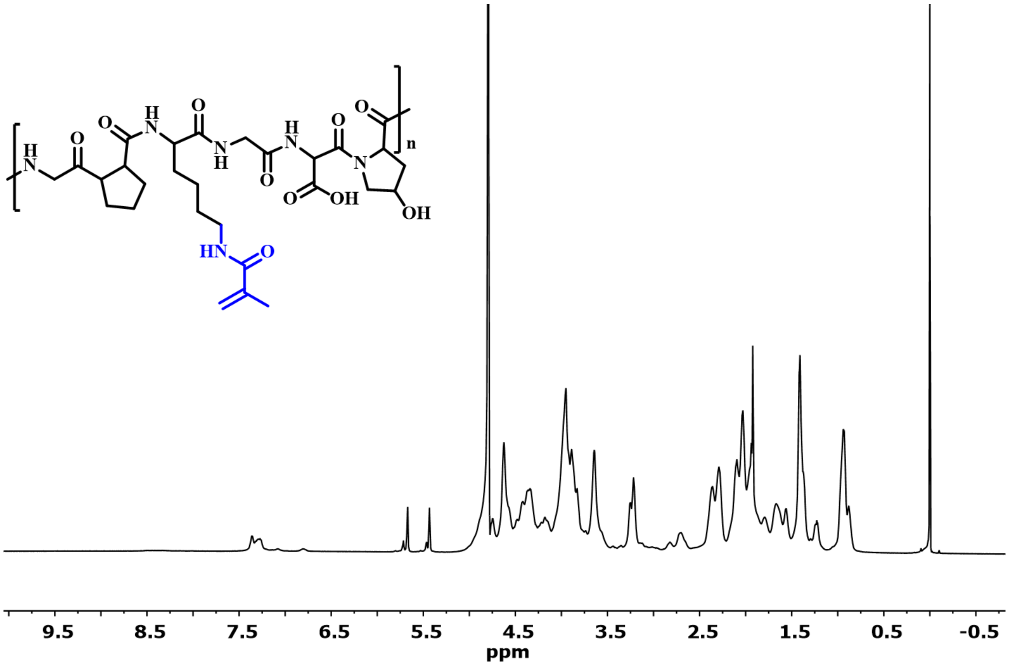


**Figure S2**. Molecular structure and $^1$H NMR ($D_2O$, 500 MHz) of GelMA.

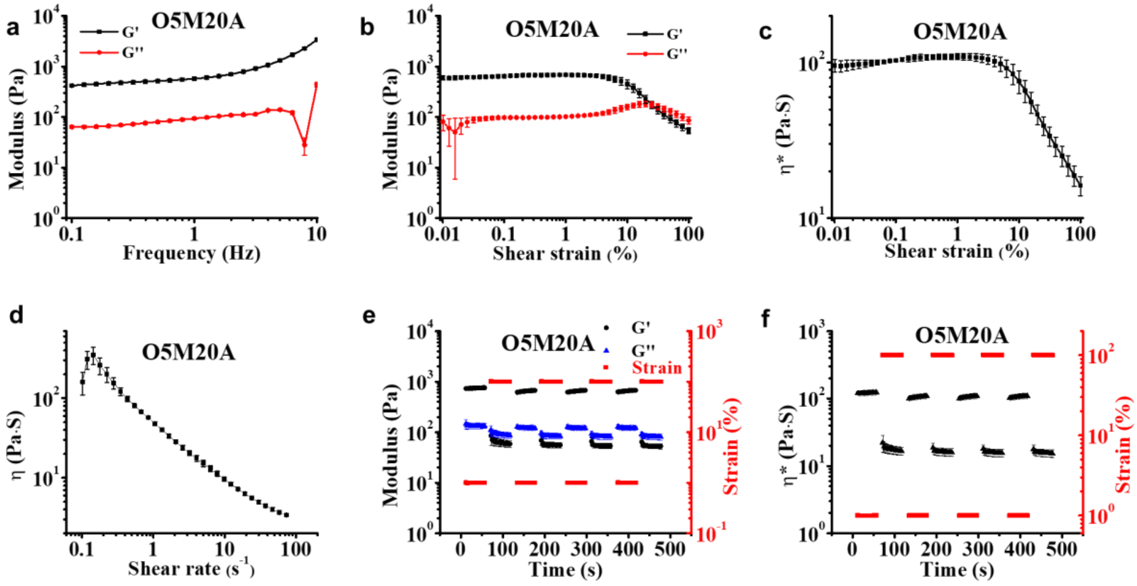


**Figure S3**. **Rheological properties of the OMA bioink.** (a) Storage modulus (G') and loss modulus (G") as a function of frequency. (b) Changes in G' and G" with increasing shear strain. (c) Complex viscosity (η*) as a function of shear strain. (d) Viscosity (η) as a function of shear rate. (e) Modulus and (f) complex viscosity changes over time under cyclic shear strain of 1% and 100%.

**Table S1**. 3D printing parameters.

| | |
|---|---|
| **Needle Size** | 22 G |
| **Printing Speed** | 4 mm/s |
| **Extrusion rate** | 1.2 μL/s |
| **Infill Density** | 60% |
| **Layer Height** | 0.66mm |
| **Printing Pattern** | Rectilinear |

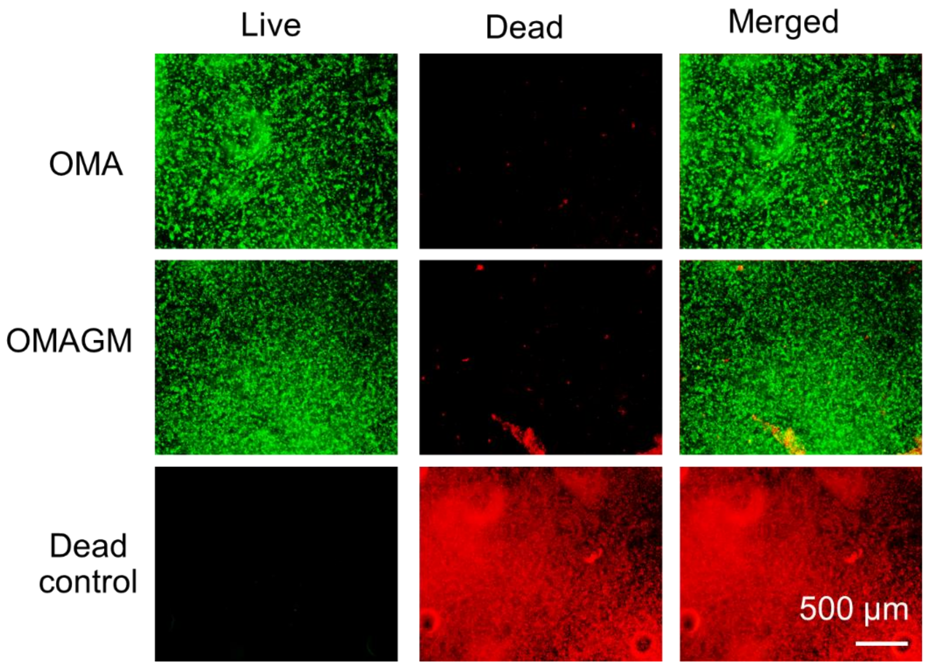


**Figure S4**. Live/dead staining images of cell-laden constructs following 4 hours of culture in cell growth medium. Conditions: cell density: 100M; UV: 15 s at 20 mW/cm$^2$; disc dimensions: $d_0$ = 8.0 mm, h = 1.0 mm.

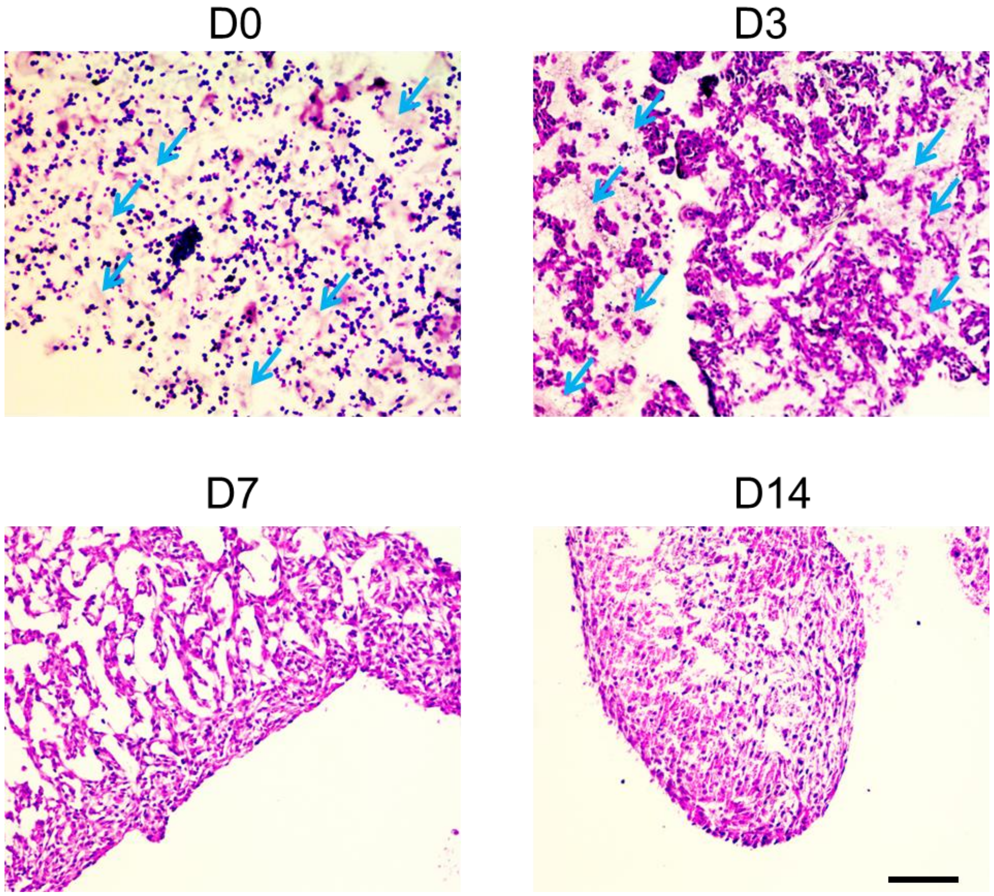


**Figure S5**. Representative H&E staining images of tissue-only constructs formed from the OMAGM group at different time points. Red arrows in the D0 and D3 images highlight a few regions of the hydrogel matrix stained in light purple color. Scale bar: 100 μm.

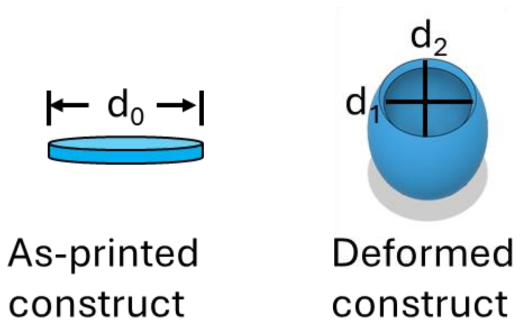


$$Contraction(\%) = \frac{(d_1+d_2)/2}{d_0} \times 100\% = \frac{d_t}{d_0} \times 100\%$$

**Figure S6**. Quantification methodology of contraction deformation in cell-laden constructs.

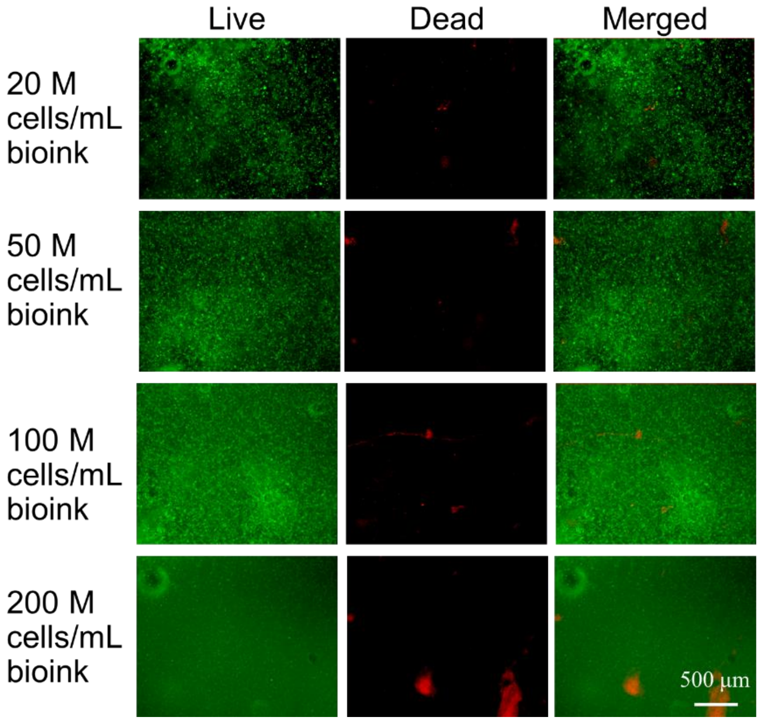


**Figure S7**. Live/dead staining images of cell-laden constructs at varying cell densities. Conditions: UV: 15 s at 20 mW/cm$^2$; disc dimensions: $d_0$ = 8.0 mm, h = 1.0 mm.

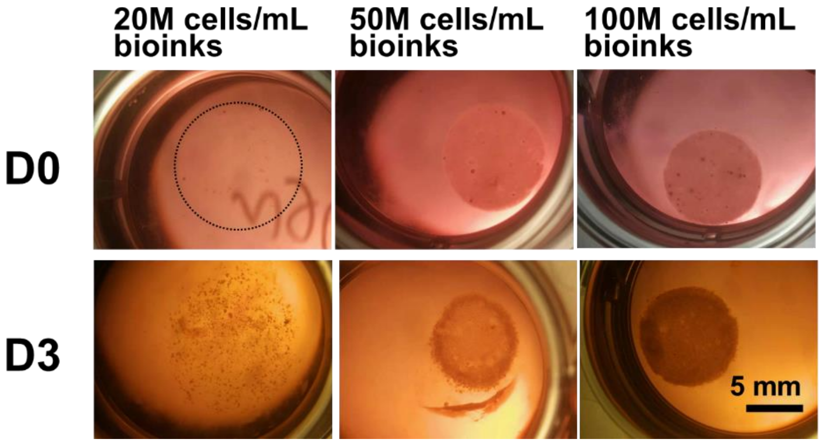


**Figure S8**. Representative images of disc constructs with varying cell densities at D0 and D3 of culture. Conditions: UV: 15 s at 20 mW/cm$^2$; disc dimensions: $d_0$ = 8.0 mm, h = 1.0 mm.

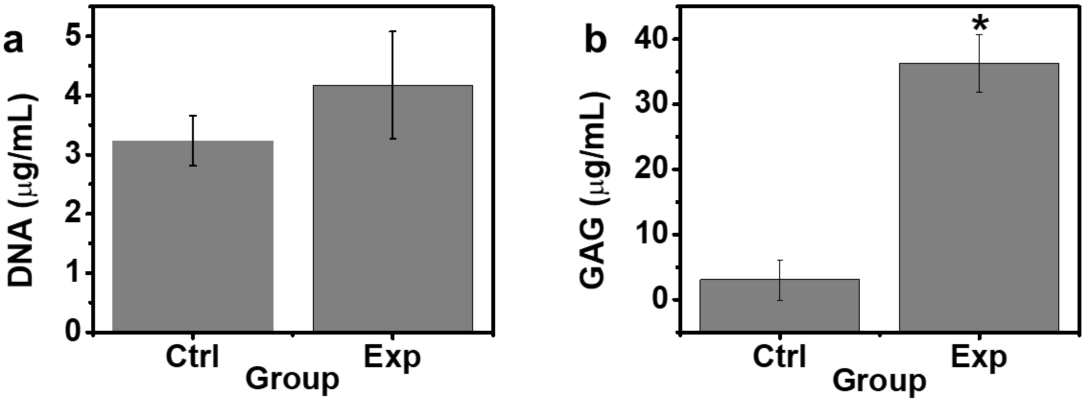


**Figure S9**. Biochemical analysis of (a) DNA and (b) GAG contents in Ctrl and Exp constructs at D14. *$p < 0.05$.

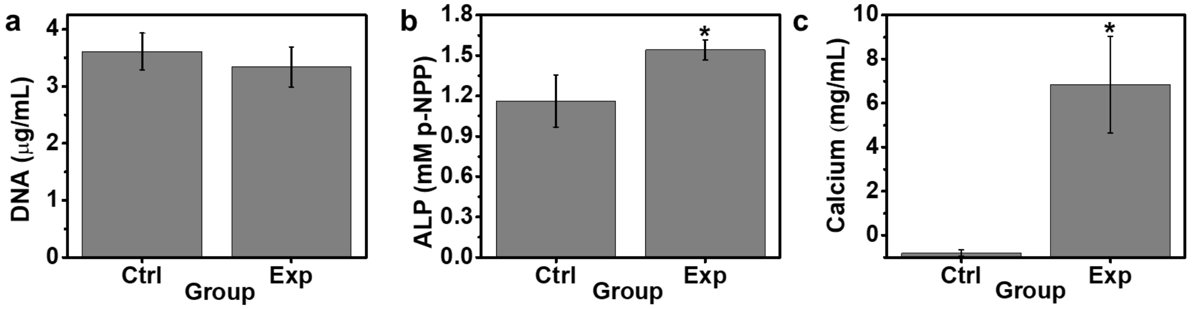


**Figure S10**. Biochemical analysis of (a) DNA, (b) ALP, and (c) calcium contents in Ctrl and Exp constructs at D14. $*p < 0.05$.
